# Contrasting population structures coexist in a strain-resolved estuarine microbiome

**DOI:** 10.64898/2026.03.20.713316

**Authors:** Lauren M. Lui, Torben N. Nielsen

## Abstract

Whether co-occurring microbial populations share similar evolutionary dynamics remains poorly understood, because strain-level resolution across many lineages simultaneously has been unachievable from metagenomic assembly. Here we apply 720 Gbp of Nanopore sequencing and the assembler myloasm to a South San Francisco Bay microbiome, recovering 488 high-quality single-contig genomes — including 328 circular chromosomes — without manual curation. Strain-level resolution revealed striking contrasts in population structure: *Pelagibacter* yielded 78 high-quality single contig genomes spanning 18 species with no two sharing >=99% average nucleotide identity (ANI) — every genome a distinct strain, consistent with phage-driven frequency-dependent selection — while HIMB114 was nearly monotypic, with 9 of 11 high-quality genomes in a single species, suggesting periodic selection or a recent sweep. Using unsupervised Markov clustering of variational autoencoder embeddings, we assigned 99.3% of contigs >=5 kbp to a taxonomic domain and resolved nearly 50% of the community to species level. The assembly also captured nearly 95,000 viral contigs (>=5 kbp) including 502 giant viruses, novel eukaryotic lineages, and mercury resistance genes reflecting the Bay’s contamination legacy. These contrasting patterns — coexisting in a single 10-liter sample — reveal that fundamentally different evolutionary strategies operate simultaneously within a complex microbiome, a level of resolution not previously demonstrated from *de novo* assembly.

We’d like to note that the results of this study mark a milestone in a 10 year hunt for Torben to get as many complete *Pelagibacter* genomes from a metagenome as possible.

## Introduction

Microbial communities routinely contain hundreds to thousands of co-occurring species across all domains of life [1, 2]. The central challenge of metagenomics is decomposition: resolving an environmental DNA sample into its constituent genomes and assigning each to a taxonomic and functional context. While short-read metagenomics has been extraordinarily productive, the fragmented assemblies it produces — typically with N50 values of 1–5 kbp — impose fundamental limits on genome recovery and strain resolution [3, 4].

Long-read sequencing, particularly from Oxford Nanopore Technologies (ONT), has transformed this landscape by producing reads long enough to span most genomic repeats, enabling the assembly of complete or near-complete genomes directly from metagenomic data [5, 6, 7]. In our original study of a South San Francisco Bay microbiome, we used 150 Gbp of Nanopore sequencing to produce an assembly of 2.95 Gbp with an N50 of 62 kbp, from which we recovered 68 near-complete genomes through extensive manual curation involving iterative read recruitment, reassembly, and quality assessment [8]. Our analysis estimated approximately 500 prokaryotic species in the sample and predicted that ∼1 Tbp of sequencing would be required for full community decomposition.

Two developments since the original study have improved the prospects for comprehensive metagenome decomposition. First, myloasm, a metagenome assembler that achieves strain-level resolution through polymorphic k-mers, has substantially improved contiguity and completeness. myloasm’s high-resolution string graph construction produces 3-fold more complete circular contigs than competing assemblers while maintaining low duplication rates [9]. Second, we recently introduced an unsupervised clustering pipeline based on variational autoencoder (VAE) embedding of k-mer frequencies, k-nearest-neighbor (k-NN) graph construction, and Markov clustering (MCL) [10]. This allows for taxonomic classification at the domain level for nearly all contigs and a large portion to the species level.

A fundamental but largely untested ecological question is whether co-occurring microbial populations exhibit similar or contrasting diversity patterns. Frequency-dependent selection by lytic phages is predicted to maintain high strain diversity in abundant host populations [11, 12], but this prediction has been difficult to test across multiple lineages in the same community because previous assemblies lacked the resolution to distinguish closely related strains. We explore this question using 720 Gbp of Nanopore sequencing — nearly five times our original depth — from the same South San Francisco Bay sample, combined with myloasm and unsupervised VAE/MCL clustering. This combination makes it possible, for the first time, to compare population structures across dozens of co-occurring genera at strain-level resolution from a single *de novo* metagenome assembly. Here we discuss (1) the improvements of the metagenome assembly (for bacteria, archaea, viruses, plasmids, and picoeukaryotes) after using these methodological advances and additional sequencing, (2) analysis of contrasting population structures enabled by the improved assembly, and (3) the results of the VAE/MCL clustering for taxonomic assignment.

## Methods

### Sample collection and DNA extraction

We collected a 10-liter surface water sample from USGS Station 36 in the South San Francisco Bay in July 2022. DNA was extracted using bead-based methods and filtered through 0.1 µm polyethersulfone membranes. Library preparation included size selection for fragments >10 kbp. Full details are described in Lui & Nielsen (2024) [8].

### Nanopore sequencing

Metagenome sequencing was performed on six Oxford Nanopore Technologies PromethION flow cells using R10.4.1 chemistry (Supplementary Table S1). Overall mean read length was 7,275 bp (N50 range 10.7–14.9 kbp; longest read 1.62 Mbp). Data from flow cells X0216 and X0220 were used in the original study [8]; the remaining four are new. Two MinION flow cells (R9.4.1 chemistry) used in the original study were excluded from this analysis because R9.4.1 reads have substantially higher error rates than R10.4.1, which could compromise assembly quality.

### Basecalling and quality control

Raw signal data were basecalled using Dorado 0.9.6 (Oxford Nanopore Technologies) with SUP (super-accuracy) models: dna_r10.4.1_e8.2_400bps_sup@v4.1.0 for LBNL_8_36 and LBNL_26_36, and dna_r10.4.1_e8.2_400bps_sup@v5.0.0 for X0244 and X0248. X0216 and X0220 used the original basecalls described in [8] and were not re-basecalled. Per-flow-cell read statistics (Supplementary Table S1) were generated using NanoPlot 1.46.2. No additional read filtering or adapter trimming was applied prior to assembly; myloasm performs internal read selection.

### Metagenome Assembly

Per-contig sequencing depth was estimated by myloasm from read coverage during assembly graph construction. All reads were assembled with myloasm 0.4.0 [9] using default parameters and 50 threads. Assembly required 912 hours of wall-clock time with 1,872% CPU utilization and a peak memory of 1.17 TB. The primary assembly (assembly_primary.fa) was used for all downstream analyses. Alternate contigs identified by myloasm were excluded from the primary analysis. To disentangle the effects of assembler choice and sequencing depth, a subset assembly was performed using only the two LBNL flow cells (265 Gbp) with the same myloasm version and parameters (Supplementary Table S3). The original assembly from [8] used Flye 2.9.1.

### Gene prediction

We predicted open reading frames for all 374,926 contigs using Pyrodigal 3.6.3 [13, 14] (a Python reimplementation of Prodigal [14]) in metagenomic mode, producing GFF3-formatted annotations and 19,562,001 coding sequence (CDS) features. Coding density was calculated per contig as the fraction of nucleotide positions covered by at least one predicted CDS, using interval union to handle overlapping genes. Median coding density varied characteristically by domain: Bacteria 88.7%, Viruses 81.5%, Archaea 80.3%, and Eukaryota 54.7% (Supplementary Fig. S1). These domain-specific distributions provide independent validation of the consensus domain assignments. Viral and eukaryotic coding densities should be interpreted with caution, as Pyrodigal is optimized for prokaryotic gene prediction and may mispredict gene boundaries in both eukaryotes and Nucleocytoviricota, which share eukaryote-like codon usage and, in some families, introns.

### Taxonomic annotation

We used five independent tools for taxonomic annotation, as described in detail in [10] with the following parameters specific to this dataset:

#### GTDB-Tk

Prokaryotic taxonomy was assigned using GTDB-Tk v2.6.1 [15, 16] with the GTDB release 226 reference database on all contigs >=100 kbp (35,389 contigs). An expanded analysis was subsequently performed on all contigs >=5 kbp (328,584 contigs) to assess classification depth across the full size range.

#### SSU rRNA extraction and classification

SSU rRNA genes were identified using cmsearch from Infernal 1.1.5 [17] with Rfam 15.0 covariance models and extracted using custom Python code. Classification was performed using vsearch 2.30.4 –usearch_global against the SILVA 138.2 NR99 database [18] at a minimum identity of 80%.

#### Tiara

Domain-level classification (Bacteria, Archaea, Eukarya, organelle) was performed using Tiara 1.0.3 [19] on all contigs >=3 kbp (346,952 contigs).

#### geNomad

Viral and plasmid sequences were identified using geNomad 1.11.2 [20] with the genomad_db v1.9 database and default settings on all contigs. Circularity and topology annotations from geNomad were not used; myloasm’s circularity calls were used exclusively, as they are based on assembly graph structure rather than terminal repeat detection.

#### MMseqs2

Protein-level taxonomy was assigned using MMseqs2 (version 18.8cc5c) taxonomy [21, 22] with default sensitivity and e-value settings against the NCBI NR database (downloaded 28 February 2026) for all contigs >=100 kbp. Per-ORF classifications were aggregated to contig-level consensus using 80% majority voting at each taxonomic rank independently. Because Pyrodigal-predicted ORFs serve as input, protein-level taxonomy for Nucleocytoviricota may be less reliable than for prokaryotes.

### Antimicrobial and metal resistance screening

Antimicrobial resistance, metal resistance, and stress response genes were identified using AMRFinderPlus 4.2.7 [23] with the –plus flag (extended search including stress and virulence genes) and database version 2026-01-21.1. All predicted proteins from Pyrodigal (19.6 million CDS) were searched using 32 threads with default identity and coverage thresholds.

### Genome quality assessment

Genome completeness and contamination were assessed using CheckM2 1.1.0 [24] on all contigs >=500 kbp (3,738 contigs), treating each contig as an individual genome. High-quality genomes were defined as >=90% complete with <5% contamination, meeting the MIMAG [25] completeness and contamination thresholds (note: MIMAG high-quality draft status additionally requires the presence of 23S, 16S, and 5S rRNA genes and >=18 tRNAs, which were not assessed here).

### Eukaryotic gene content assessment

BUSCO 6.0.0 [26] was run in genome mode with the eukaryota_odb10 lineage dataset (255 markers) on all eukaryotic contigs >=500 kbp (157 contigs identified as eukaryotic in the consensus domain pipeline). Default parameters were used with miniprot 0.18-r281 as the gene predictor.

### Pairwise ANI analysis

Average nucleotide identity was computed for 12 prokaryotic genera selected to span a range of abundance and population structure (*Pelagibacter*, *Pelagibacter_A*, SAR86, HIMB114, HIMB59, *Luminiphilus*, MS024-2A, *Actinomarina*, BACL14, Casp-actino5, *Pontimonas*, UBA3478) using skani 0.3.1 [27] in triangle mode with default parameters (ANI detection threshold ∼80%; pairs below this threshold report zero). All contigs >=500 kbp assigned to each genus by GTDB-Tk were extracted as individual FASTA files and compared pairwise. Population structure analyses used only high-quality genomes (>=90% completeness, <5% contamination) to avoid artifacts from incomplete or chimeric contigs inflating single-linkage clusters. Species-level clusters were defined by single-linkage clustering at 95% ANI [28]. Single-linkage is conservative for species counting; its chaining tendency merges borderline clusters, yielding equal or fewer species than average-linkage approaches, and this risk is mitigated when comparing high-quality genomes rather than short contig fragments. For comparison with the original assembly, the 68 manually curated bins from Lui & Nielsen (2024) [8] were compared against the myloasm assembly using skani in search mode; best matches were identified by highest ANI (Supplementary Table S2 and S3).

### Rarefaction analysis

To test whether diversity contrasts among genera could be explained by unequal genome counts, we performed rarefaction analysis on the high-quality-only pairwise ANI matrices. For each sample size *k* (5 to *n* genomes), *k* genomes were drawn without replacement 1,000 times, the subsampled ANI matrix was clustered by single-linkage at 95% ANI, and species were counted. For cross-genus comparison, all genera with >=11 high-quality genomes were subsampled to n=11 (matching HIMB114, the smallest genus with a clear population pattern). Pairs with ANI below skani’s detection threshold (∼80%) were assigned a distance of 25 percentage points. All analyses used a fixed random seed (42) for reproducibility.

### Consensus taxonomy

A consensus domain assignment was generated for each contig >=5 kbp using a priority-based framework:

1. GTDB-Tk assignments (highest priority for prokaryotes with phylum-level or deeper classification)
2. SILVA SSU rRNA-based classification
3. Tiara domain-level classification
4. MMseqs2/NR consensus
5. geNomad plasmid classification (lowest priority)
6. geNomad virus classification overrides all other sources except GTDB-Tk prokaryotic assignments

GTDB-Tk calls were demoted in two situations. First, domain-only calls (classified to Bacteria or Archaea but without phylum resolution) were deferred to subsequent sources; this identified 14 contigs, of which 11 were reclassified as Eukaryota based on concordant Tiara and SILVA evidence and three remained Bacteria. Second, calls flagged by GTDB-Tk as “genome domain questionable” were cross-validated against Tiara: when Tiara independently classified such a contig as eukaryotic, the GTDB-Tk assignment was rejected. This identified 11 additional misclassified contigs, including four placed by GTDB-Tk into Asgardarchaeota — a known confound, since Asgardarchaeota share many proteins with eukaryotes and GTDB-Tk can attract divergent eukaryotic sequences to this branch of the archaeal tree. Ten were reclassified as Eukaryota and one was confirmed as Bacteria by SILVA.

Tiara “prokarya” assignments were resolved by cluster-based majority voting (724 contigs resolved) or by prior probability (remaining 2,790 assigned to Bacteria based on the 99.2% bacterial proportion among resolved prokaryotes).

### VAE embedding and clustering

Contig clustering followed the VAE/MCL pipeline described in [10]. Briefly, canonical k-mer frequency vectors (1-through 6-mers; 2,772 features) were computed for all 374,926 primary contigs from the myloasm assembly and transformed to Euclidean space via per-group centered log-ratio (CLR) transformation. The CLR-transformed vectors were projected into a 384-dimensional latent space using a VAE trained on 668,505 sequences comprising 655,640 NCBI reference replicons (February 2026) and 12,865 marine eukaryote reference sequences from 16 genomes spanning picoeukaryotes, diatoms, protists, fish, and marine mammals (NCBI_euk_5 model; 1,000 training epochs; best validation loss 85.33 at epoch 971).

For visualization, 2D t-SNE projections were computed using openTSNE with perplexity 30 and cosine distance metric (Fig. 2). A k-nearest-neighbor graph was constructed with k=15, maximum Euclidean distance 5.0, inverse-distance edge weighting (w = 1/(d + 0.1)), and maximum in-degree 100, yielding 130,959 connected nodes and 1,032,971 edges. MCL 22-282 clustering at inflation I=3.0 (selected from I in {1.4, 2.0, 3.0, 4.0, 5.0, 6.0} by GC span and cluster size) produced 21,448 multi-member clusters and 9,387 singletons.

### Taxonomy propagation

Contigs within MCL clusters that lacked GTDB-Tk classification inherited the consensus taxonomy of their cluster members, provided >=80% of classified members agreed. This propagated phylum-level taxonomy to 14,724 previously unclassified contigs.

## Results

### Improvements in metagenome assembly driven primarily by strain-resolving assembler

Our original study estimated ∼1 Tbp for full decomposition of the South San Francisco Bay microbiome [8]. To achieve this goal we needed not only more sequencing but also a metagenome assembler that could assemble data at that scale. We sequenced more of the original DNA from a single surface water sample from USGS Station 36 in the South San Francisco Bay (collected July 2022) resulting in data from six ONT PromethION flow cells, generating 720 Gbp (Supplementary Table S1). Flye was used for the original assembly; it is known to have input limits at about half a terabase and thus was no longer suitable. We chose to use myloasm because not only could it handle the amount of data, but it resolves closely related co-occurring strains as separate contigs rather than collapsing them into consensus sequences, enabling the strain-level population analyses described below.

Assembly with myloasm produced 374,926 primary contigs totaling 20.5 Gbp, with an N50 of 89,830 bp and a largest contig of 9.28 Mbp (Supplementary Table S3) — a 7-fold increase in assembly size over our original Flye assembly [8]. With this expansion, the N50 improved from 62 to 90 kbp, reflecting deeper sequencing and myloasm’s greater contiguity (a subset assembly at 265 Gbp achieved an intermediate N50 of 75 kbp; Supplementary Table S3). myloasm identified 6,161 contigs as circular and an additional 10,832 as possibly circular. Of the 374,926 contigs, 3,738 were >=500 kbp — a reasonable lower bound for the length of a complete prokaryotic chromosome — totaling 4.45 Gbp.

The ∼1 Tbp goal for sequencing was based on an estimate of ∼500 bacterial and archaeal species and relative abundances, so we next sought to evaluate the number of prokaryotic genomes recovered. CheckM2 [24] assessment of the 3,738 contigs >=500 kbp identified 488 as high-quality single-contig genomes (>=90% complete, <5% contamination). Of these, 328 (67%) were circular, suggesting complete chromosome recovery. Pairwise ANI analysis confirmed minimal redundancy: of the 488, only 11 pairs exceeded 99% ANI, all representing co-occurring strains within the same species, and no pair exceeded 99.8% ANI. The recovery of these high-quality genomes represents a 7-fold improvement over the 68 near-complete genomes recovered from our original assembly, which required intensive manual curation [8].

To directly assess the improvement in high-quality genome recovery, we compared the 68 original bacterial bins to the myloasm assembly using skani [27] (Supplementary Table S3). All 68 were recovered at >=95% ANI, with every best-matching myloasm contig being circular. Of the 68, 48 (71%) were recovered as high-quality genomes without any manual curation. The remaining 20 bins were not recovered as single high-quality genomes, primarily because myloasm resolved their source populations into many individual strain-level contigs rather than collapsing them into single consensus sequences — for example, where the original assembly produced one contaminated *Opacimonas* bin (100% completeness, 21% contamination), myloasm resolved two clean high-quality genomes of the same species (99.1% ANI) — one circular — plus fragments from lower-abundance strains. Among the 23 original bins with >=10% contamination — where multiple organisms had been merged into a single bin — 22 were resolved into clean single contigs (<5% contamination). Three *Luminiphilus* bins — two heavily contaminated (100%/97% and 95%/64%) — were resolved into 14 high-quality genomes spanning 11 species, eight of them circular; similarly, two of six Pseudohongiellaceae bins were heavily contaminated (100%/90% and 100%/86%) and were resolved into eight high-quality genomes across four species, five of them circular. In both cases, what appeared to be single contaminated genomes were amalgams of multiple co-occurring species that we could not separate with our previous methods.

With 720 Gbp — approximately 72% of the target 1 Tbp and among the deepest long-read metagenome assemblies to date [29, 30] — the 7-fold improvement in assembly size (20.5 Gbp vs 2.95 Gbp) and the recovery of high-quality genomes (488 vs 68) reflects both deeper sequencing and myloasm’s strain-resolution capabilities [9]. To disentangle these two factors, we assembled a subset of two flow cells (265 Gbp, ∼1.8× the original depth) with myloasm (Supplementary Table S4). The original Flye assembly produced only 13 high-quality single-contig genomes directly; the 68 reported in [8] required additional read recruitment, Canu re-assembly, and manual curation. The myloasm subset yielded 244 high-quality genomes including 173 circular — an 18.8-fold improvement over the comparable Flye baseline (13 automated high-quality genomes) despite only 1.8-fold more data. Increasing the data to 720 Gbp (2.7-fold) doubled the count to 488. Cross-assembly ANI comparisons confirm high concordance between assemblies (Supplementary Fig. S2; Supplementary Table S2). The disproportionate gain from switching assemblers versus adding depth indicates that the strain-resolving assembler is the primary driver of improvement, with sequencing depth providing further gains (though read quality also differs between flow cells; Supplementary Table S1).

### Strain recovery reveals taxonomic novelty and discordance of genome ANI and SSU sequence identity

Of the 488 high-quality bacterial and archaeal genomes, 184 (38%) were likely novel species: none received a Genome Taxonomy Database (GTDB) species assignment, and 91 had measurable ANI to a reference genome, all below the 95% species boundary [28, 31]; the remaining 93 fell below skani’s detection limit (∼80% ANI), indicating no closely related reference genome exists (Supplementary Table S5). Among the 91 with measurable ANI, the most divergent had values of 83–85% to their nearest GTDB references — well below the 95% species threshold — yet several were complete, circular chromosomes with near-perfect quality metrics. These include an *Opacimonas* sp. genome (99.7% complete, 0% contamination, circular, 2.9 Mbp; 83.2% ANI), a *Puniceispirillum*/SAR116 genome (100% complete, 0.3% contamination, circular, 2.4 Mbp; 84.1% ANI), and an RS24/Parvibaculales genome (100% complete, 0% contamination, circular, 2.0 Mbp; 84.4% ANI).

Eleven genomes were classified only to family level, representing candidate novel genera from deeply branching lineages. These candidate novel genera span eight families across seven phyla: a circular 896 kbp Babelota (previously Dependentiae) genome (Chromulinivoraceae) — a phylum whose characterized members are obligate intracellular parasites of protists [32, 33] — with only 87.4% small subunit (SSU) rRNA identity to its nearest SILVA match; a Margulisbacteria genome from a deeply branching lineage sister to Cyanobacteria [34]; a Nanobdellota (Woesearchaeales) archaeon; two Patescibacteria (Gracilibacteria); two Oligoflexia predatory bacteria (Bdellovibrionota_B); a Francisellaceae; a Schleiferiaceae (Bacteroidota); and two Rickettsiaceae — obligate intracellular parasites assembled directly from bulk environmental DNA, one matching *Candidatus Cryptoprodotis* (a parasite of ciliates) at 93.6% SSU identity.

Several of the most divergent genomes exhibited a striking discordance between SSU rRNA identity and whole-genome ANI. The *Opacimonas* genome had 99.9% SSU identity to its nearest reference but only 83.2% ANI, and the SAR116 genome had 99.6% SSU identity but 84.1% ANI. This pattern of conserved ribosomal markers masking extensive genome divergence is consistent with previous observations in cultured isolates [35, 36] and SAR11 [37], but those earlier reports relied on cultivated strains or fragmentary metagenome-assembled genomes (MAGs). Here, the recovery of complete, circular chromosomes with >15 percentage points of SSU-ANI discordance directly from *de novo* metagenomic assembly demonstrates that this phenomenon can now be captured at genome-complete resolution, and illustrates a practical limitation of 16S-based surveys: highly conserved SSU sequences can both underestimate the genomic novelty present in a community and misidentify organisms by assigning them to taxa with near-identical 16S but deeply divergent genomes.

### Analysis of community composition indicates improved genome assembly across viruses and all domains of life

We applied five independent annotation tools to characterize the assembled contigs: GTDB-Tk [15, 16] for prokaryotic taxonomy, SILVA/vsearch [18] for SSU rRNA-based classification, Tiara [19] for domain-level eukaryote/prokaryote discrimination, geNomad [20] for virus and plasmid identification, and MMseqs2 [21, 22] against the NCBI non-redundant (NR) protein database for protein-level taxonomy (see Methods). Results were integrated into a consensus taxonomy using a priority-based framework, with additional resolution provided by cluster-based taxonomy propagation (described below in “Compositional clustering validates community structure”). The domain-level counts reported here reflect this full consensus pipeline.

Of the 328,584 contigs >=5 kbp, 326,277 (99.3%) were assigned to a taxonomic domain (Fig. 1a). The community was dominated by Bacteria (194,889 contigs; 59.3%), followed by viruses (94,954; 28.9%), Eukaryota (34,764; 10.6%), Archaea (1,629; 0.5%), and 41 geNomad-classified plasmids not assigned to a host domain. Only 2,307 contigs (0.7%) remained unassigned at the domain level. We extracted 8,450 SSU rRNA gene sequences from 7,476 contigs — a 16-fold increase over the 524 recovered in the original assembly.

**Figure 1.**
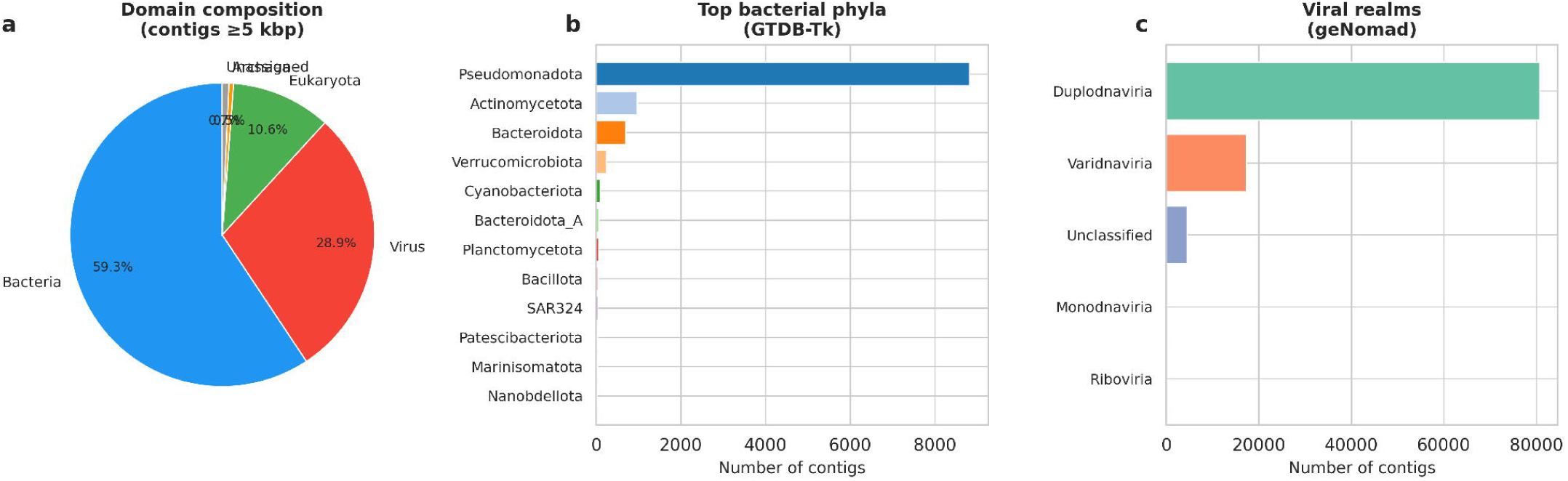
Community composition of the South San Francisco Bay microbiome. (a) Domain-level assignment of 328,584 contigs >=5 kbp, showing Bacteria (59.3%), Virus (28.9%), Eukaryota (10.6%), Archaea (0.5%), and unassigned (0.7%). We excluded contigs shorter than 5kb (46,342, ∼12% of all contigs) from this analysis because the taxonomic classifications are generally not reliable. (b) Top bacterial phyla by contig count (GTDB-Tk classification). (c) Viral realm composition (geNomad classification).

**Figure 2.**
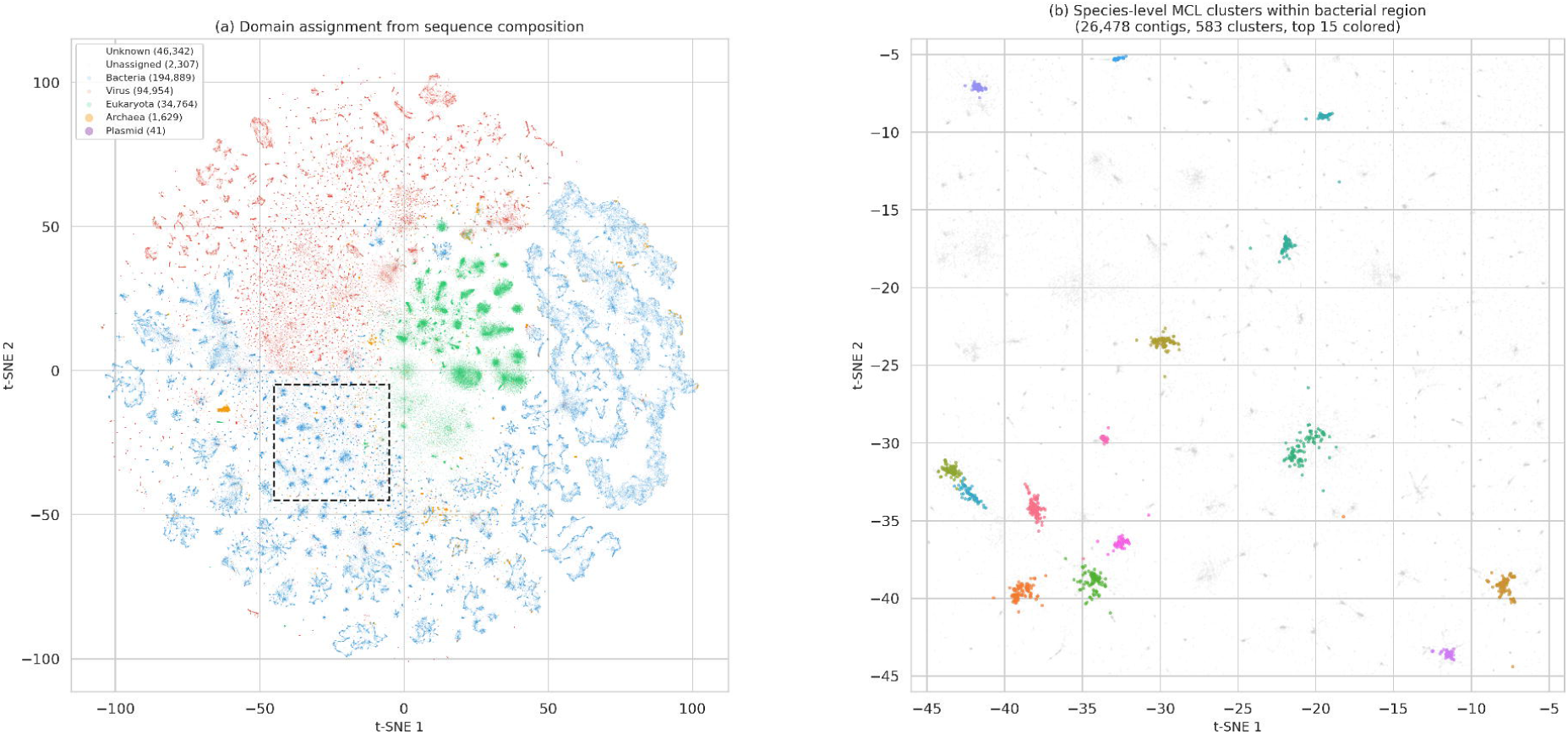
VAE embedding captures community structure at domain and species resolution. (a) t-SNE projection of the VAE latent space for all 374,926 assembled contigs, colored by consensus domain assignment (Bacteria blue, Virus red, Eukaryota green, Archaea orange). The embedding separates taxonomic domains purely from oligonucleotide composition, without taxonomy input. Dashed box indicates the bacterial region shown in (b). (b) Zoom into the boxed region, with contigs colored by MCL cluster membership (top 15 clusters by size). Each tight cluster corresponds to a species-level grouping; 583 clusters are present in this region alone, illustrating the species-level resolution achieved by unsupervised clustering.

Classification against the SILVA 138.2 NR99 database using vsearch assigned 8,225 (97.3%) to known taxa at a mean identity of 99.2%. The SSU-based taxonomy showed 99.1% concordance with Tiara domain-level classifications, validating both independent approaches.

#### Improvements in prokaryotic genome assembly include 78 Pelagibacter and others rarely recovered from metagenomes

GTDB-Tk classified 8,565 prokaryotic contigs >=100 kbp to at least the phylum level (8,555 after cross-validation demotion; see Methods), with Pseudomonadota (6,724) dominating (Fig. 1b), followed by Actinomycetota (770), Bacteroidota *sensu lato* (653, including Bacteroidota_A), and Verrucomicrobiota (205). At the species level, Pelagibacterales (SAR11) were the most abundant, contributing 4,148 contigs >=100 kbp across five genera, multiple species, and hundreds of strains — consistent with their status as the most abundant bacterioplankton lineage in the ocean [38]. Among these, 78 high-quality *Pelagibacter* genomes span 18 species by 95% ANI clustering, with no two genomes sharing >=99% ANI — each representing a distinct strain (see “Strain resolution and population structure” below). SAR86 clade Gammaproteobacteria [39, 40], which dominated our original analysis at 10.3% of reads, were again prominent.

Beyond the expected marine microbiome constituents, the assembly yielded categories of organisms rarely recovered as high-quality or complete genomes from metagenomes. The seven largest circular contigs (5.1–6.4 Mbp) were all complete Planctomycetes–Verrucomicrobiota–Chlamydiae (PVC) superphylum genomes (Planctomycetes and Verrucomicrobiota [41, 42]) — genomes typically fragmented in metagenomic assemblies. The most abundantly sequenced organism was not a typical marine dominant but *Opacimonas* (Alteromonadaceae, Gammaproteobacteria), with 256 classified contigs at a mean sequencing depth of 9,974× — approximately 93-fold higher than the assembly-wide mean of 107×. Despite the extreme depth, only two were recovered as high-quality genomes (one circular), likely because the high strain diversity fragments the assembly.

Most (178/256) are novel species by GTDB-Tk classification. *Opacimonas* is a recently described genus with only two cultured species, both from diatom phycospheres [43]; its extreme abundance in a South San Francisco Bay summer sample is unexpected and may reflect a bloom event at the time of sampling.

Archaeal representation improved based on the number of full length SSUs detected and high-quality genomes recovered. Of the full length SSUs, 22 were classified as archaea, in comparison to only one recovered in the original assembly. Among the 280 archaeal contigs retained after cross-validation (from 286 GTDB-Tk classifications; see Methods), notable lineages included DPANN (Diapherotrites–Parvarchaeota–Aenigmarchaeota–Nanoarchaeota–Nanohaloarchaeota) archaea (Nanobdellota, 4 contigs including a high-quality 1.22 Mbp Woesearchaeales genome at 93% completeness — notable as DPANN archaea are rarely assembled from metagenomes due to their low abundance), Marine Group II Thermoplasmatota (3 contigs), and an ammonia-oxidizing Thermoproteota (*Nitrosopumilus*). Previously no high-quality archaeal genomes were recovered and only fragments indicated the presence of multiple archaeal taxa. The recovery of the archaeal SSUs and Nanobdellota genome likely reflect improvements due to deeper sequencing.

#### Notable improvements in giant virus genome recovery

Although the proportion of viral contigs decreased compared to the original assembly (28.9% vs 42%), this likely reflects greater contiguity based on the number of recovered circular genomes (5539 vs 3112), including circular jumbo phages and giant viral genomes (89 vs 14). The larger number of circular viral genomes indicates we have more complete viral genomes; we do not know the true improvement of complete dsDNA viral genomes in the sample since many are likely linear. geNomad identified 102,506 viral contigs across all sizes (2.2-fold more than the 45,788 in our original analysis), of which 94,954 (>=5 kbp) were retained in the consensus domain analysis. Among these, 8,897 exceeded 100 kbp and 502 exceeded 300 kbp — the latter a commonly used size threshold for jumbo phages and giant viruses (Nucleocytoviricota) [44, 45] — a 5.7-fold increase over our original count.

The viral community was dominated by Duplodnaviria (75,452 contigs; 79%; Fig. 1c), consistent with the well-documented dominance of tailed bacteriophages in aquatic metagenomes [46]. Varidnaviria accounted for 16,583 contigs, with Algavirales (Phycodnaviridae) dominant within this realm, reflecting the presence of their algal hosts in the estuary.

While the 502 jumbo phages and giant viruses (>=300 kbp) noted above include only the largest members, the full Nucleocytoviricota (NCLDV) assemblage is far more extensive: 16,711 contigs totaling 1.12 Gbp — 5.5% of the entire assembly. Of these, 35 exceeded 1 Mbp and 3 exceeded 2 Mbp, with the largest at 2.76 Mbp (the original assembly only had 3 viral contigs >1 Mbp). Six circular virus-classified contigs >=500 kbp represent putatively complete giant virus genomes, each encoding multiple major capsid protein genes and other NCLDV hallmark genes; three additional circular contigs >=500 kbp carry small (58–73 kbp) NCLDV provirus insertions within bacterial chromosomes. The family-level diversity by geNomad classification (which uses legacy taxonomy that may not fully reflect recent International Committee on Taxonomy of Viruses (ICTV) revisions to Nucleocytoviricota [47]) spanned Phycodnaviridae (13,746), Mimiviridae (413), Poxviridae (67), Iridoviridae (30), Schizomimiviridae (20), Marseilleviridae (13), Asfarviridae (13), and ∼2,400 unclassified; the Poxviridae and Iridoviridae assignments are unexpected in a marine sample and may reflect misclassification of divergent marine NCLDV that fall outside well-characterized families. Additionally, 403 Preplasmiviricota contigs were recovered — a phylum that includes virophages (Lavidaviridae) and Polinton-like viruses. The co-occurrence of algal hosts, their NCLDV parasites, and Preplasmiviricota in a single sample is consistent with, but does not by itself demonstrate, a multi-trophic interaction network [48, 49, 50].

#### Plasmids and mobile elements are the main repositories for antimicrobial and metal resistance genes

geNomad identified 4,269 plasmid contigs, of which 210 carried conjugation genes (F-type, T-type, and MOB mobilization systems) and 320 harbored antimicrobial resistance (AMR) genes by geNomad’s functional annotation (which uses a broader definition than curated AMR databases; see below).

Screening all 19.6 million predicted proteins with AMRFinderPlus [23] identified 109 antimicrobial resistance genes and 156 metal/stress resistance genes across 228 contigs. The metal resistance signal was dominated by mercury (116 genes on 93 contigs) and arsenic (39 genes on 39 contigs). Mercury resistance genes were frequently organized in intact *mer* operons [51] (20 contigs carried >=2 *mer* genes). Of the 32 plasmids carrying any AMRFinderPlus resistance hit, 26 (81%) carried mercury resistance genes, making mercury detoxification the dominant resistance function on plasmids in this community. Arsenic resistance showed a distinct taxonomic distribution, concentrated in Verrucomicrobiota (5 contigs carrying *arsD* metallochaperone genes), with one carrier at 17,817× sequencing depth — the most abundant resistance-carrying contig in the assembly.

Antimicrobial resistance was modest (109 genes). Beta-lactam resistance accounted for the majority (77 hits), predominantly class B3 metallo-beta-lactamases likely representing intrinsic rather than acquired resistance, followed by lincosamide (14), fosfomycin (7), phenicol (3), and other minor classes. Only 10 of the 488 high-quality prokaryotic genomes carried any resistance gene (7 antimicrobial, 3 metal/stress only), indicating low prevalence among the dominant community members.

#### Eukaryotic fraction expanded

The relative fraction of contigs classified as eukaryotic increased compared to the original assembly (18.3% vs 4%) and largely recapitulated the taxonomic classifications (Fig 1a). Tiara classified 63,323 of 346,952 contigs as eukaryotic, with an additional 792 plastid and 369 mitochondrial sequences classified separately as organellar. SSU rRNA analysis identified 160 eukaryotic sequences across seven major lineages: Chlorophyta (62 SSUs), Cryptophyceae (30), Stramenopiles (23), Cercozoa (19), Opisthokonta (10), Ciliophora (8), Dinoflagellata (7), and one unresolved eukaryote. MMseqs2 protein-level taxonomy of the 892 eukaryotic contigs >=100 kbp identified Chlorophyta as the dominant phylum (225 contigs; *Picochlorum* 80, *Ostreococcus* 72), followed by Bacillariophyta (122; *Skeletonema* 31), while 545 contigs (61%) received no phylum-level assignment, suggesting substantial uncharacterized eukaryotic diversity. Six SSU sequences fell below 90% identity to the nearest SILVA reference, spanning four supergroups: Archaeplastida (Chlorophyta, 81.9%, nearest *Picochlorum*), SAR (Cercozoa 82.3% and 86.2%; Stramenopiles 82.9%, nearest Chrysophyceae), Cryptista (Cryptophyceae, 86.8%), and Obazoa (Opisthokonta, 86.9%, nearest Cryptomycota/LKM11), suggesting novel lineages across the eukaryotic tree.

Benchmarking Universal Single-Copy Orthologs (BUSCO) [26] assessment of the 157 eukaryotic contigs >=500 kbp recovered 243 of 255 (95.3%) eukaryota_odb10 markers across the combined set, with 83.1% duplicated — indicating that the eukaryotic community as a whole carries near-complete gene content, distributed across multiple species and strains. Individual contigs carry up to 49 markers (19%), consistent with chromosome-scale fragments of picoeukaryotic genomes (*Ostreococcus* and *Picochlorum* genomes are 13–15 Mbp distributed across ∼20 chromosomes).

### Compositional clustering validates community structure

#### Assessment of VAE embedding and graph-based clustering for taxonomic assignment

To improve taxonomic assignment of contigs, we applied the VAE/MCL clustering pipeline described in Lui and Nielsen [10] to group contigs by oligonucleotide composition similarity. Briefly, k-mer frequency profiles were computed for all 374,926 contigs and projected into a reduced feature space using a variational autoencoder (VAE) trained on 668,505 reference sequences (655,640 NCBI reference replicons and 12,865 marine eukaryote genomes) (see Methods for full parameters). From the embeddings, a k-nearest-neighbor graph connected 130,959 contigs through 1,032,971 edges. MCL clustering [52, 53] produced 21,448 multi-member clusters containing 121,572 contigs, plus 9,387 singleton clusters. A t-distributed stochastic neighbor embedding (t-SNE) projection of the VAE embedding (Fig. 2a) showed that the learned representation naturally separated taxonomic domains without any taxonomy input — Bacteria, Virus, Eukaryota, and Archaea each occupy distinct regions of the latent space. Within any one domain, further structure is apparent: zooming into a single bacterial quadrant (Fig. 2b) reveals hundreds of tight, well-separated clusters, each corresponding to a species-level grouping.

Full assessment of the VAE/MCL pipeline for taxonomic coherence is described in (VAE paper), but we also assessed cluster quality here using GC content span. GC content is a strong proxy for taxonomic coherence. If highly similar taxa are assigned to the same cluster, we would expect the GC span to be narrow. For this San Francisco Bay microbiome, the median GC span was 0.6 percentage points (pp), with a 90th percentile of 1.8 pp, indicating compositionally coherent clusters (Fig. 3a). Domain-level purity was 99.7%: of 21,415 clusters with >=2 domain-assigned members, 21,342 had all members in the same domain (Fig. 3b). The 73 mixed-domain clusters were predominantly Bacteria/Virus pairs (53 clusters, likely prophage-containing contigs) and Eukaryota/Virus pairs (17, likely giant virus integration events), with 3 clusters containing other domain combinations (Bacteria/Eukaryota, Archaea/Bacteria). Phylum-level purity among the 1,009 clusters with >=2 GTDB-Tk phylum calls was 99.9%, with a single mixed cluster (Bacteroidota/Pseudomonadota, size 2).

**Figure 3.**
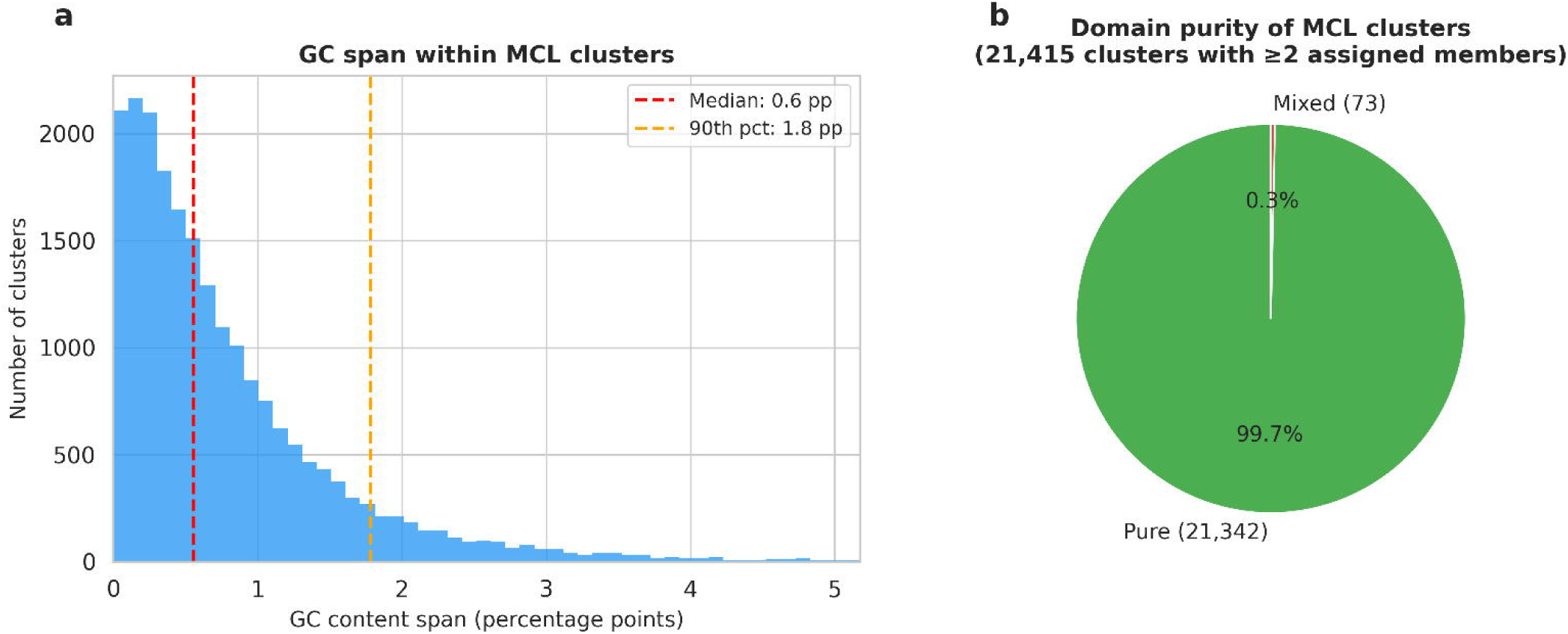
Cluster quality assessment. (a) Distribution of GC content span (percentage points) across MCL clusters at I=3.0. Median span 0.6 pp, 90th percentile 1.8 pp. (b) Domain-level purity across 21,415 clusters with >=2 domain-assigned members, showing 99.7% with uniform domain assignment.

#### Taxonomy propagation

Cluster membership enabled taxonomy propagation: contigs lacking direct GTDB-Tk classification inherited the consensus taxonomy of their cluster, provided >=80% of classified members agreed. This step assigned phylum-level taxonomy to 14,724 contigs that had no prior sub-domain classification. An additional 724 Tiara “prokarya” (unresolved bacteria/archaea) assignments were resolved to Bacteria (700) or Archaea (24) based on cluster majority, with the remaining 2,790 assigned to Bacteria by prior probability (99.2% of resolved prokaryotes are bacterial).

#### Plasmid–host co-clustering

Narrow-host-range plasmids are known to ameliorate their oligonucleotide composition toward that of their hosts [54, 55], though many retain distinct compositions [56], so a k-mer-based clustering pipeline should group ameliorated plasmids with their host genomes. Of the 4,269 plasmid contigs, 461 (10.8%) co-clustered with cellular hosts in 233 mixed clusters. Of the 36 clusters containing two or more GTDB-Tk-classified host contigs, all had hosts belonging to a single genus — zero cross-genus or cross-phylum mixing. The best example was a *Luminiphilus* cluster containing 62 host contigs — including four high-quality genomes — alongside four plasmids (56–122 kbp). No plasmids co-clustered with archaeal or eukaryotic contigs. The genus-level coherence across all classifiable clusters is consistent with the VAE embedding capturing the compositional signature of plasmid–host amelioration [54, 55], though independent validation of these putative host assignments remains to be performed.

### Large differences population structure co-exist exemplified by 12 abundant genera, including *Pelagibacter*

Pairwise ANI analysis (skani [27]) of the high-quality genomes from 12 abundant genera selected to span a range of population structures revealed striking differences in population structure within a single environmental sample (Fig. 4). We investigating intra-genus and inter-genus diversity, and used rarefaction analysis to account for sequencing depth.

**Figure 4.**
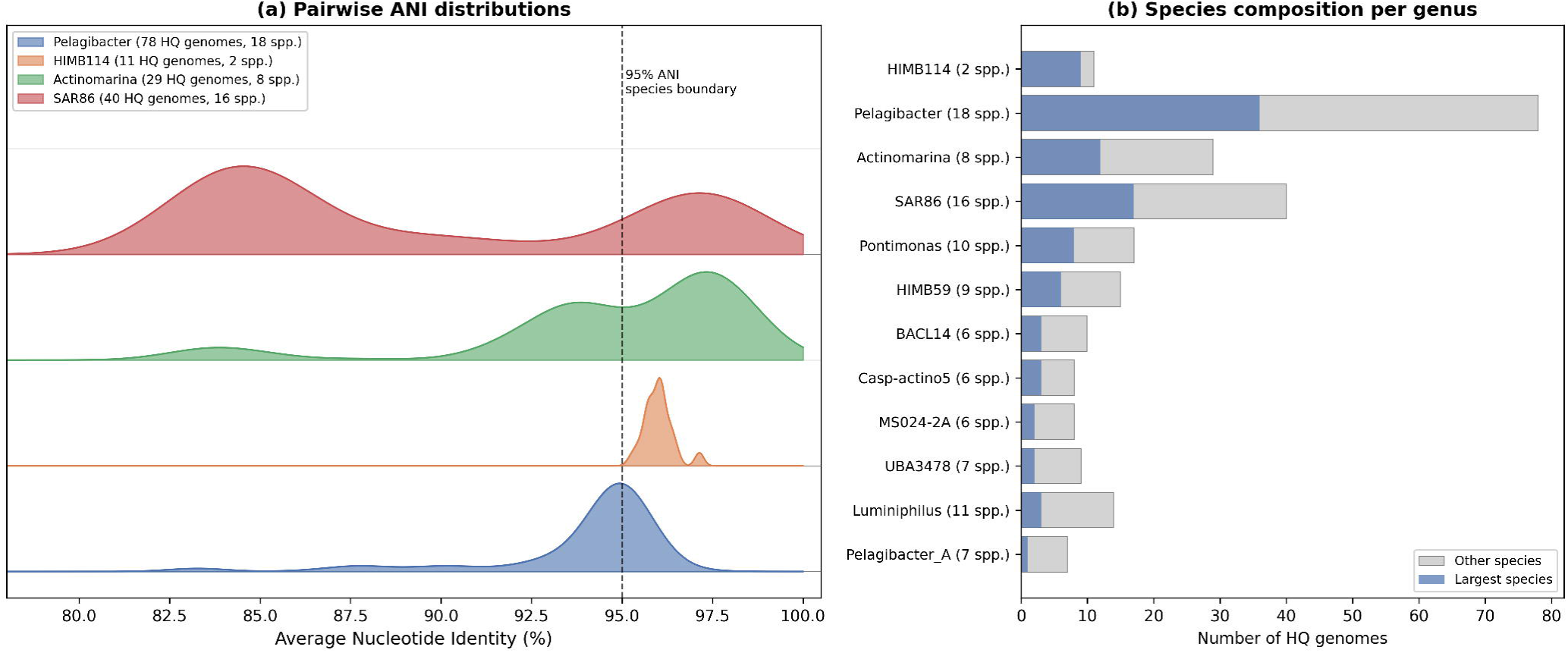
Population structure across 12 abundant genera. (a) Pairwise ANI distributions for five genera illustrating contrasting population structures: *Pelagibacter* (78 high-quality genomes, 18 species), HIMB114 (11 high-quality genomes, nearly monotypic), *Actinomarina* (29 high-quality genomes, 8 species), SAR86 (40 high-quality genomes, 16 species), and *Luminiphilus* (14 high-quality genomes, maximally species-diverse). All genera use high-quality genomes only (>=90% completeness, <5% contamination) to avoid artifacts from incomplete contigs. Dashed line indicates the 95% ANI species boundary. (b) High-quality genome and species counts for all 12 genera, showing the fraction in the largest species (blue) versus remaining species (gray).

The *Pelagibacter* assemblage showed the greatest intra-genus diversity. Restricting the analysis to the 78 high-quality genomes to exclude chimeric and incomplete contigs, pairwise ANI resolved 18 species by single-linkage 95% ANI clustering (Fig. 5). The two largest species by GTDB-Tk classification are sp008638165 (36 genomes) and sp028229795 (26 genomes), with 16 unclassified genomes forming smaller clusters or singletons — eight of these circular. No two genomes share >=99% ANI (maximum 97.6%), meaning every genome represents a distinct strain. The VAE/MCL clustering independently validated this strain-level diversity. Among the 4,148 Pelagibacterales contigs >=100 kbp, 4,127 were placed into 1,549 MCL clusters with near-perfect genus-level segregation (98–100% same-genus preference). Within *Pelagibacter* sp008638165, 2,124 contigs were distributed across 1,024 clusters, each strain’s slightly different k-mer signature placing it in a distinct neighborhood of the VAE latent space.

**Figure 5.**
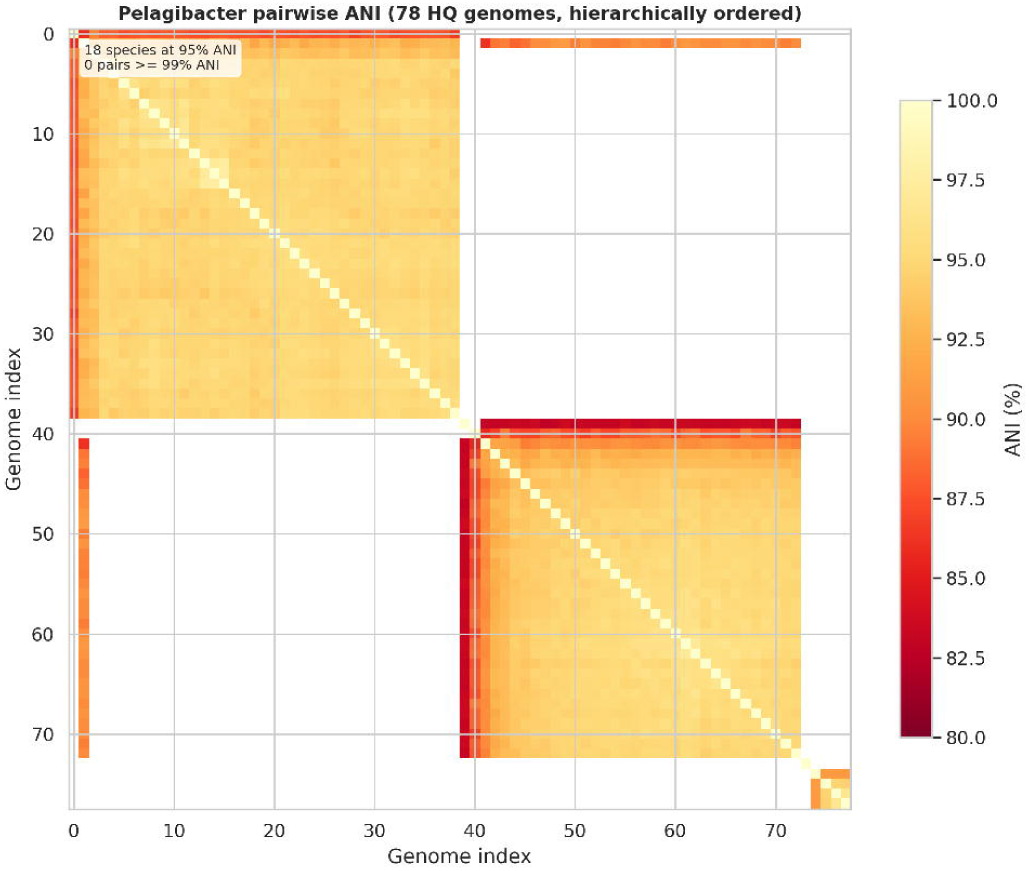
*Pelagibacter* strain diversity. Pairwise ANI heatmap of 78 high-quality *Pelagibacter* genomes (>=90% complete, <5% contamination), hierarchically ordered by average-linkage clustering. Two dominant species blocks are visible: sp008638165 (upper left, 36 genomes) and sp028229795 (lower right, 26 genomes), with 16 unclassified genomes forming singletons or small clusters at the margins. White cells indicate pairs below skani’s detection threshold (∼80% ANI). No two genomes share >=99% ANI (maximum 97.6%), indicating that every genome represents a distinct strain.

Other genera showed markedly different patterns. HIMB114 was nearly monotypic: 9 of 11 high-quality genomes fell within a single species (median ANI 96.0%). In contrast, *Luminiphilus* was maximally diverse, with 14 high-quality genomes distributed across 11 species and no cluster larger than 3. SAR86 showed moderate concentration: 40 high-quality genomes resolved into 16 species, with one dominant species of 17 genomes.

MS024-2A, *Actinomarina*, *Pontimonas*, and HIMB59 all showed high species diversity relative to sample size (6–10 species from 8–29 high-quality genomes). Among these 12 genera, no pair of high-quality genomes shared >=99% ANI. Pseudohongiellaceae illustrated the practical impact of strain resolution: the original Flye assembly produced 6 bins for this family, two heavily contaminated (90% and 86%), while myloasm resolved the same population into 5 complete circular genomes (2.3–4.2 Mbp, all >=97% complete, <1% contamination) and 98 total contigs spanning multiple strain variants.

Rarefaction analysis supported these diversity contrasts as robust to unequal sampling. If one genus is highly abundant and another is not, if the sequencing is too low it may not capture the true diversity of the rarer genus. Thus we conducted rarefaction analysis to test if these diversity contrasts were artifacts of sequencing depth. When *Pelagibacter* was subsampled to n=11 high-quality genomes (matching HIMB114), it retained a median of 5 species (95% CI: 3–9) compared to HIMB114’s two (Fig. 6). Notably, despite the reputation of *Pelagibacter* of having a large number of species compared to others in the marine environment, *Luminiphilus*, SAR86, HIMB59, and *Pontimonas*, were more diverse after accounting for sequencing depth (Fig 6b).

**Figure 6.**
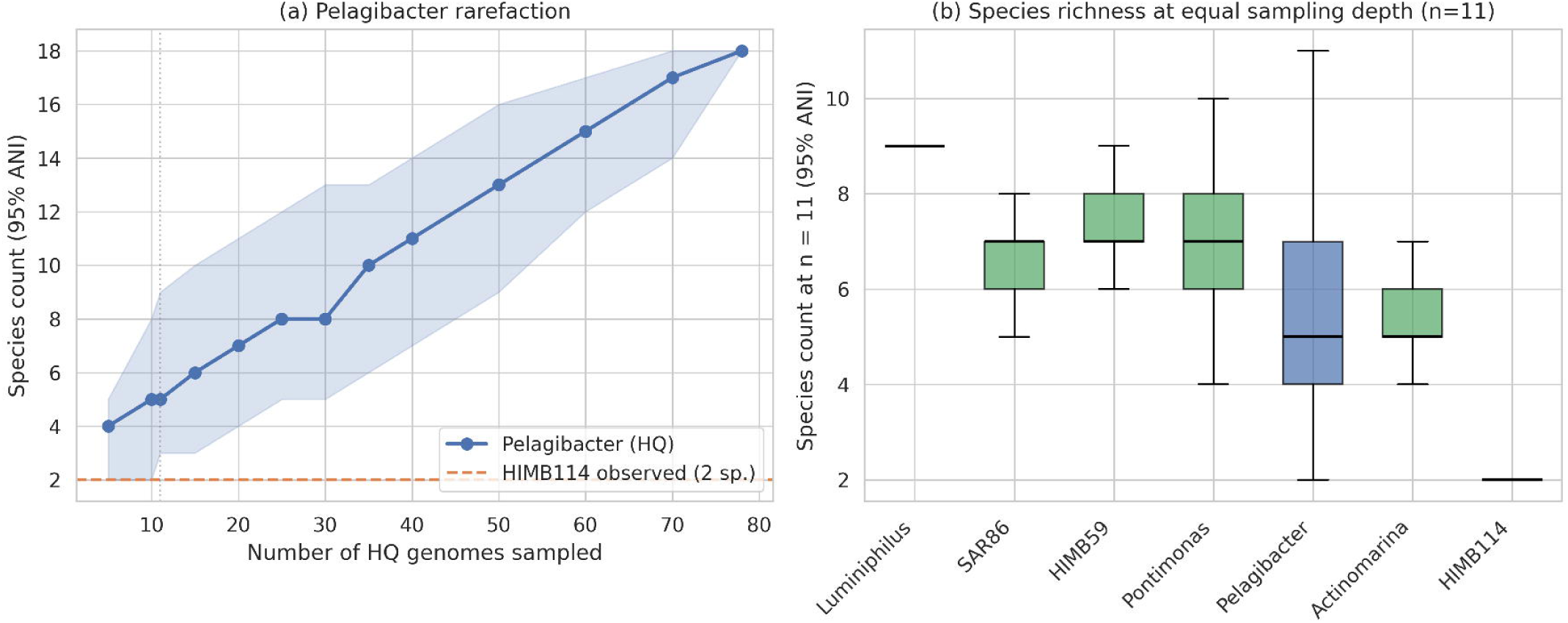
Species richness rarefaction analysis. (a) Rarefaction curve for *Pelagibacter* high-quality genomes (n=78), showing median species count (95% ANI single-linkage clustering) as a function of the number of high-quality genomes subsampled (1,000 replicates per sample size; shaded area: 95% CI). Dashed orange line indicates HIMB114’s observed species count (2 species at n=11). Even at n=5, *Pelagibacter* exceeds HIMB114’s diversity; the curve shows no sign of saturating at n=78. (b) Cross-genus comparison at equal sampling depth (n=11). All genera with >=11 high-quality genomes were subsampled to n=11 (1,000 replicates). HIMB114 (2 species) is an outlier at the low end; *Luminiphilus*, HIMB59, and *Pontimonas* are the most species-rich. The diversity contrasts among genera persist after controlling for sampling effort.

### Taxonomic depth is affected by genomic mass and coding density signatures

While 99.3% of contigs >=5 kbp received domain-level classification, the depth of taxonomic assignment varied markedly with contig length (Fig. 7). Only 33,538 contigs (10.2%) had phylum-level or deeper taxonomy, reflecting the sharp decline in marker gene recovery below ∼100 kbp. However, when weighted by genomic mass (contig length x sequencing depth), 72.9% of the community was classified to phylum or better, and 49.6% to species level. This disparity arises because the longest, most deeply sequenced contigs— which represent the most abundant organisms — are also the most likely to receive deep taxonomic annotation. Contigs >=100 kbp accounted for 77.3% of total genomic mass, and among these, 67–99% (depending on length bin) received phylum-level classification or better. Extending GTDB-Tk to all 328,584 contigs >=5 kbp confirmed this pattern: of the 293,195 contigs in the 5–100 kbp range, only 2,542 (0.9%) received phylum-level or deeper classification, compared to 24.2% of contigs >=100 kbp.

**Figure 7.**
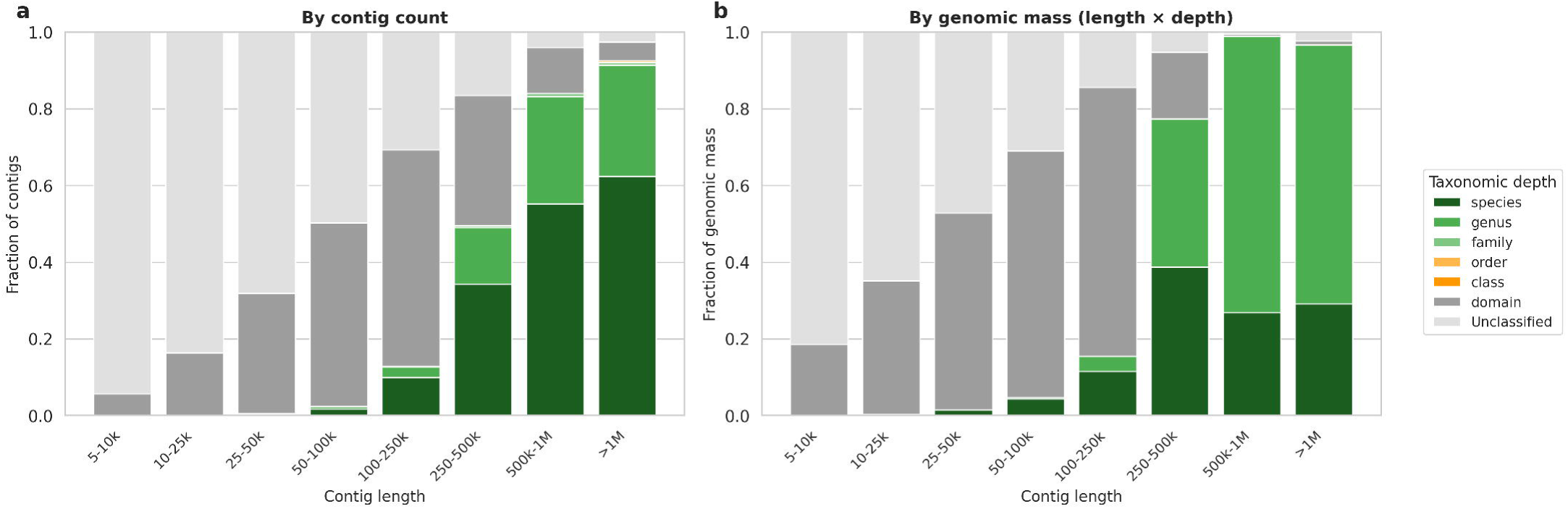
Taxonomic depth as a function of contig length. Stacked bar charts showing the fraction of contigs classified to each taxonomic depth (domain, phylum, class, order, family, genus, species) by contig length bin, weighted by (a) contig count and (b) genomic mass (length x depth). The sharp transition at ∼100 kbp reflects the practical limit of marker gene-based classification on shorter contigs.

## Discussion

### Strain resolution and population structure

The contrasting population structures described above — from the nearly monotypic HIMB114 to the hyper-diverse *Pelagibacter* (Fig. 4) — coexist in a single 10-liter water sample. The *Pelagibacter* pattern, in which no two of 78 high-quality genomes share>=99% ANI, is consistent with frequency-dependent selection driven by lytic phage predation: the hypervariable regions bounded by rRNA genes in *Pelagibacter* genomes [37] have been suggested as possible targets of phage recognition. Although high rates of homologous recombination within SAR11 [37] could blur ANI-based species boundaries, the discrete clustering observed here — with no genome pairs between 95% and 99% ANI — suggests that recombination has not erased species-level differentiation in this assemblage. While kill-the-winner dynamics [11] predict species-level coexistence, the within-species strain diversity is consistent with constant-diversity dynamics [12], in which phage recognition of strain-specific surface structures prevents any single genotype from dominating — a mechanism for which experimental evidence exists in *Prochlorococcus* [57]. The HIMB114 pattern — 9 of 11 genomes in a single species — is consistent with periodic selection, a recent selective sweep, or a founder effect from recent colonization; distinguishing among these scenarios will require temporal sampling.

*Luminiphilus*, with 11 species among 14 genomes, indicates species-level niche partitioning. No previous metagenomic assembly — including our original Flye assembly — resolved individual *Pelagibacter* genomes; as of March 2026, only 31 complete (circular) *Candidatus Pelagibacter* genomes existed in NCBI [58, 59], and our 78 high-quality genomes — of which 31 are circular — substantially expand available genomic diversity. Because deeper sequencing enables detection of lower-abundance strains, the 18 species observed may underestimate true diversity; however, neutral evolutionary processes cannot be excluded without temporal replication.

Single-cell genomics has previously revealed hundreds of coexisting *Prochlorococcus* subpopulations [60], and theoretical frameworks for interpreting microbial population structure from sequence data have been developed [61], but the simultaneous resolution of contrasting population structures across multiple co-occurring genera from a single *de novo* assembly is, to our knowledge, unprecedented. These patterns represent a summer snapshot; seasonal shifts may substantially alter both community composition and within-species diversity, and disentangling ecological drivers from other confounds will require multi-site and temporal comparisons.

### Mercury and metal resistance as environmental signal

The dominance of mercury resistance genes (116 of 265 resistance hits) is notable in the context of South San Francisco Bay’s well-documented mercury contamination from New Almaden mine drainage and industrial discharge [62, 63]. The prevalence of plasmid-encoded *mer* operons — 26 of 32 resistance-carrying plasmids bore mercury resistance genes — is consistent with horizontal transfer of mercury detoxification capacity within the community, possibly driven by selective pressure from bioavailable mercury. The 39 arsenic resistance genes may similarly reflect the Bay’s industrial contamination history. These results illustrate how functional gene surveys of metagenomes can reveal community-level adaptations to local environmental stressors, complementing recent genome-resolved characterizations of SF Bay sediment communities [64].

### Giant viruses and eukaryotic diversity

The 502 giant virus contigs (>=300 kbp) represent a 5.7-fold increase over our original analysis, consistent with the growing recognition of giant viruses as major components of aquatic ecosystems [44, 45, 47, 65]. The eukaryotic fraction (10.6% by the consensus pipeline, 18.3% by Tiara alone) substantially exceeds the 4% estimated originally, likely reflecting the increased assembly size and Tiara’s superior sensitivity [19]. The six deeply divergent SSU sequences spanning four supergroups at ∼82–87% identity, together with the 61% of eukaryotic contigs >=100 kbp lacking phylum-level classification, suggest substantial uncharacterized diversity in marine protist communities.

### Taxonomic novelty and its implications

That 184 of 488 high-quality genomes (38%) represent novel species — organisms with no genome in public databases above 95% ANI — adds substantially to what is known about estuarine microbial diversity. Many of these lineages would be detectable as operational taxonomic units by 16S amplicon surveys, but a 16S sequence alone reveals little about metabolic capability and, as the SSU-ANI discordances reported here illustrate, can misidentify organisms by assigning them to taxa with near-identical ribosomal markers but deeply divergent genomes. Having complete or high-quality genomes for these novel species — including obligate intracellular parasites (Babelota, Rickettsiaceae), deeply divergent chlorophytes, and poorly characterized genera such as *Opacimonas* — provides the substrate for metabolic reconstruction and hypothesis-driven cultivation. The 488 strain-level genomes also open a path to understanding *why* these organisms coexist: comparative genomics across strains within and between species can identify niche-differentiating genes and test whether functional redundancy or complementarity maintains the observed diversity.

### Unsupervised clustering versus per-genome manual curation

The 488 high-quality single-contig genomes reported here — comparable in scale to the 1,083 high-quality metagenome-assembled genomes (MAGs) recovered across 23 activated sludge plants [66], though from a single sample — required no per-genome intervention, compared to the 5–10 person-hours per genome in our original study [8]. All 68 original bins were recovered, with 48 as high-quality circular genomes and 22 of 23 originally contaminated bins resolved into clean genomes (Supplementary Table S2). The VAE/MCL pipeline [10], combined with myloasm’s contiguity, makes per-genome curation largely unnecessary. The 99.7% domain-level purity and 99.9% phylum-level purity of our MCL clusters validate this unsupervised approach at scale: the original validation [10] used our 2.95 Gbp Flye assembly, and here the same approach scales to 20.5 Gbp with 374,926 contigs while maintaining equivalent purity. The few impure clusters (73 mixed-domain, predominantly Bacteria/Virus and Eukaryota/Virus pairs) likely represent biological reality — prophage insertions and giant virus integration events — rather than methodological failure.

### Remaining challenges for metagenome decomposition

Whether decomposition is “complete” depends on the definition: 99.3% of contigs >=5 kbp are domain-assigned, but 89.1% remain at domain level only, and the long tail of rare species means complete recovery is unlikely at any practical depth. Approximately 169,000 bacterial contigs in the 5–100 kbp range lack sub-domain taxonomy because short contigs rarely contain sufficient marker genes, and the ∼198,000 contigs >=5 kbp not connected in the k-NN graph represent “dark matter” accessible through alternative approaches. Broad-host-range plasmids may act as bridge nodes in the k-NN graph, causing spurious cluster merges [67, 68]. Genome completeness estimates rely on CheckM2 marker gene sets, and assembly errors — including chimeric contigs and strain-level mosaics — cannot be excluded without independent validation. The VAE embedding model was trained on known reference organisms, which may reduce sensitivity for deeply divergent lineages.

Although rarefaction analysis shows that genus-level diversity contrasts persist at equal sampling depth (Fig. 6), within-genus diversity estimates may still be influenced by coverage differences. Finally, these results represent a single summer time point; seasonal variation remains to be characterized.

The combination of deep long-read sequencing, a strain-resolving assembler, and unsupervised clustering reveals that fundamentally different population structures — from hyper-diverse to nearly monotypic — coexist within a single microbial community. From a single 10-liter water sample, we recovered 488 high-quality single-contig genomes including 184 novel species, assigned 99.3% of contigs to a taxonomic domain, and resolved nearly half the community’s genomic mass to species level. This level of population-genetic resolution — distinguishing dozens of strains within a species from a bulk environmental sample — has not previously been achievable from *de novo* metagenomic assembly, and shifts the bottleneck from data generation and per-genome curation to biological interpretation of the contrasting evolutionary dynamics now visible within complex microbiomes.

## Data availability

Sequencing data are available at NCBI under BioProject PRJNA1435553. The assembled contigs, annotation files, and analysis notebooks will be made available when the manuscript is published.

## Code availability

All custom scripts used in this analysis will be made available on Github when the manuscript is published. The VAE/MCL pipeline is described and distributed as part of [10].

## Acknowledgements

We thank USGS for collecting samples for us, especially Erica Nejad and the crew of USGS R/V David H. Peterson. We cannot overemphasize the value of the support we have received for this project from USGS. We also thank Miten Jain for lively discussions about nanopore sequencing and providing support for the PromethION sequencing.

## Author contributions

**L.M.L.:** Conceptualization, Methodology, Investigation, Formal analysis, Data curation, Writing — original draft, Writing — review & editing. **T.N.N.:** Conceptualization, Methodology, Software, Formal analysis, Data curation, Visualization, Writing — original draft, Writing — review & editing.

## Ethics declarations

No specific permits were required for surface water collection at USGS Station 36, a publicly accessible environmental monitoring site in the South San Francisco Bay. No human or animal subjects were involved.

## Abbreviations

AMR: antimicrobial resistance
ANI: average nucleotide identity
BUSCO: Benchmarking Universal Single-Copy Orthologs
CDS: coding sequence
CLR: centered log-ratio
DPANN: Diapherotrites–Parvarchaeota–Aenigmarchaeota–Nanoarchaeota–Nanohaloarchaeota
GFF: General Feature Format
GTDB: Genome Taxonomy Database
ICTV: International Committee on Taxonomy of Viruses
k-NN: k-nearest neighbor
MAG: metagenome-assembled genome
MCL: Markov clustering
MIMAG: minimum information about a metagenome-assembled genome
NCLDV: nucleocytoplasmic large DNA virus
NR: non-redundant (NCBI protein database)
ONT: Oxford Nanopore Technologies
PVC: Planctomycetes–Verrucomicrobiota–Chlamydiae
SRA: Sequence Read Archive
SSU: small subunit (ribosomal RNA)
VAE: variational autoencoder.

## Supplementary Materials

**Supplementary Table S1. Sequencing summary.** Six PromethION flow cells generated 720.6 Gbp from 99.1 million reads.

**Supplementary Table S2. Cross-assembly genome concordance.** Each high-quality genome (>=90% completeness, <5% contamination) from the query assembly was compared to high-quality genomes in the reference assembly using skani v0.3.1 (>=80% ANI threshold). “Flye direct” = 13 high-quality single-contig genomes from the original Flye assembly (150 Gbp); “Original bins” = 68 manually curated genomes from Flye + Canu re-assembly [8]; “myloasm 265 Gbp” = 173 circular high-quality genomes from the subset assembly. Reference sets: Flye direct and myloasm 265 Gbp were compared against 328 circular high-quality genomes from the full myloasm assembly; Original bins were compared against all 488 high-quality genomes. All 13 Flye direct genomes matched a myloasm genome at >=99.2% ANI. Of the 173 subset circular high-quality genomes, 93 (54%) matched at >=99% ANI, and of the 27 without a circular high-quality match, 2 matched non-circular high-quality genomes and the remaining 25 matched non-high-quality contigs at >=98.3% ANI — in 17 cases the full assembly recovered only a fragment (median 26% of the subset genome size) and in 8 cases the assembler produced chimeric contigs merging multiple strains (22–93% contamination). Of the 68 original bins, 52 (76%) matched a myloasm high-quality genome; the 16 without a match had elevated strain variant counts (median 9 vs 5 for matched bins), suggesting that high strain diversity hinders single-contig genome recovery.

**Supplementary Table S3. Comparison of 68 original bins to myloasm assembly.** For each of the 68 near-complete genomes from the original Flye-based analysis [8], this table lists the original bin composition (number of contigs, size, completeness, contamination), the best-matching myloasm contig (completeness, contamination, length, circularity, ANI), the number of strain variants resolved by myloasm, and the GTDB genus assignment.

**Supplementary Table S4. Assembly comparison.** To disentangle the contributions of assembler choice and sequencing depth, a subset of two flow cells (LBNL_26_36 and LBNL_8_36; 265 Gbp) was assembled independently with myloasm. Annotation pipelines (SSU extraction, geNomad, domain assignment) were not run on the subset assembly. The original Flye assembly produced only 13 high-quality single-contig genomes directly; the 68 reported in [8] required additional read recruitment, Canu re-assembly, and manual curation. Cross-assembly concordance is shown in Supplementary Fig. S2.

**Supplementary Table S5. Quality metrics and GTDB-tk classification for the 488 high-quality genomes.** Genome completeness and contamination values were obtained from running CheckM2.

**Supplementary Fig. S1. Coding density by domain.** Violin plots of coding density (fraction of contig length covered by predicted CDS) for Bacteria (median 88.7%), Virus (81.5%), Archaea (80.3%), and Eukaryota (54.7%). Viral coding densities should be interpreted with caution, as Pyrodigal is optimized for prokaryotic gene prediction.

**Supplementary Fig. S2. Cross-assembly genome concordance.** (A) Best-hit ANI for each query genome against its closest match in the full myloasm assembly (720 Gbp). Each point represents one genome; unmatched genomes (no hit at >=80% ANI) are annotated. Flye direct: 13 high-quality single-contig genomes from the original Flye assembly compared to 328 circular high-quality myloasm genomes. Original bins: 68 curated genomes from [8] compared to all 488 high-quality myloasm genomes. myloasm 265 Gbp: 173 circular high-quality genomes from the subset assembly compared to 328 circular high-quality myloasm genomes. (B) Proportion of query genomes in each ANI category. Numbers inside bars indicate genome counts.

