## Supplementary figures and images for "Contrasting population structures coexist in a strain-resolved estuarine microbiome"

### Supplementary Figure 1

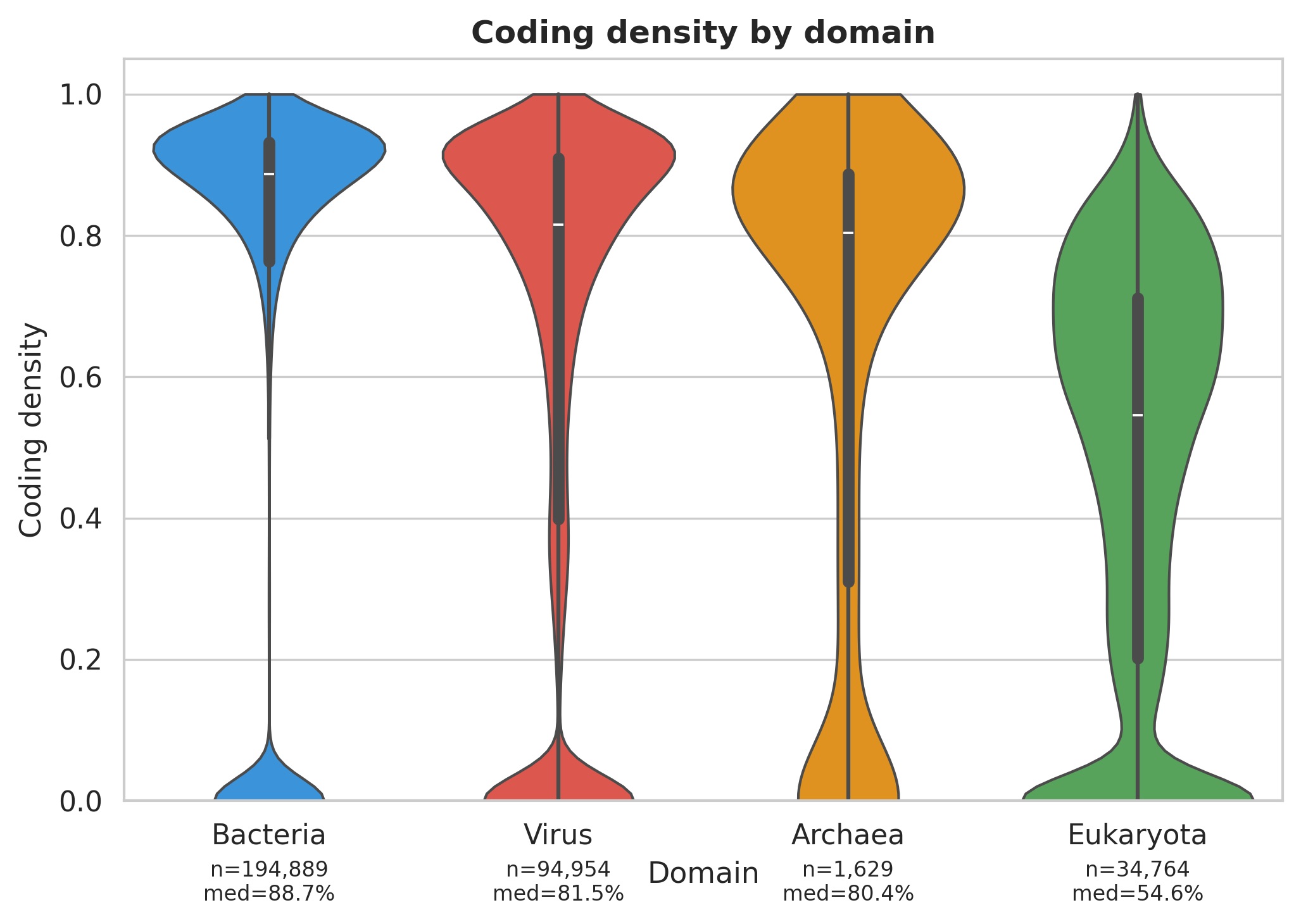

### Supplementary Figure 2

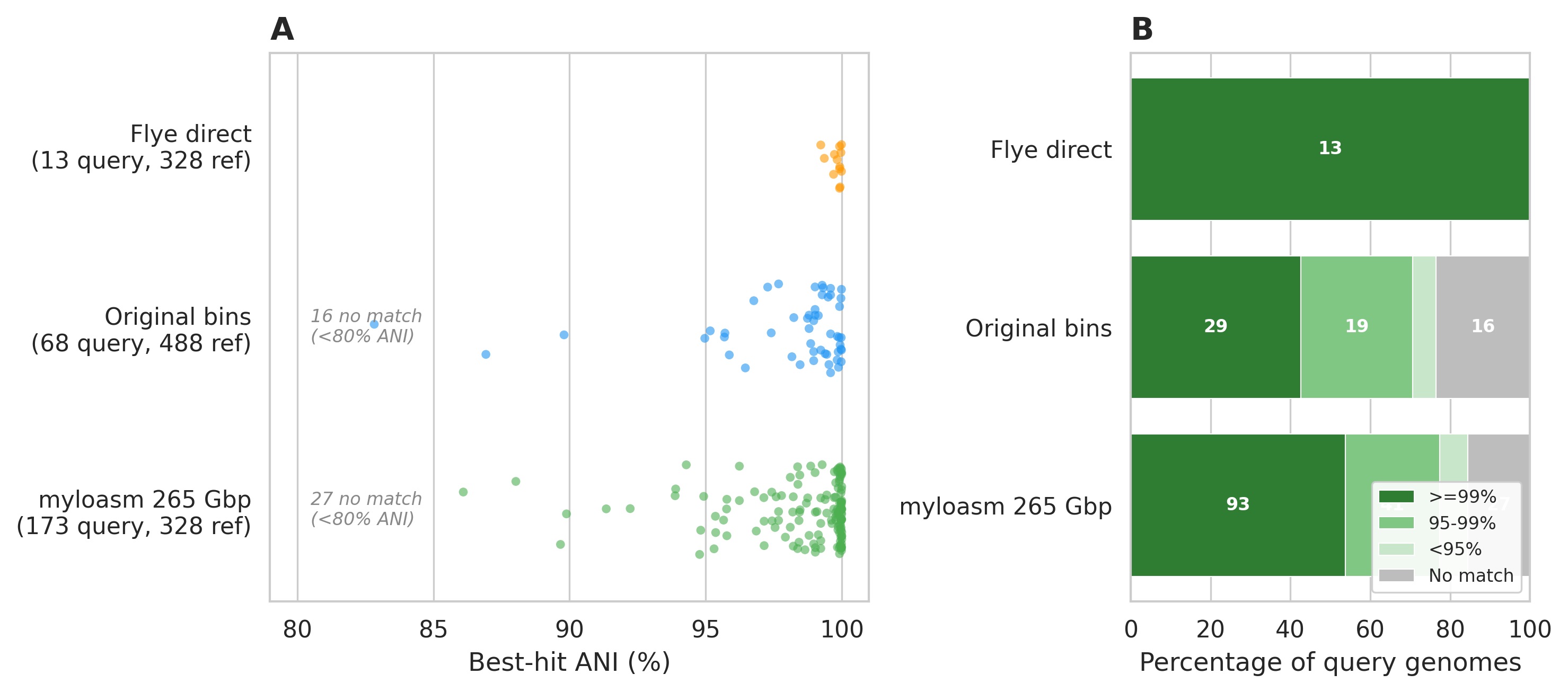
